# Aminoglycoside antibiotics drive RNA phase separation

**DOI:** 10.64898/2026.08.07.743409

**Authors:** Julian von Hofe, Mechi Chen, Christine Choi, Ka Wing Leung, Aarav Kokane, Kfir B. Steinbuch, Deyuan Cong, Moeka Sasazawa, Yitzhak Tor, Saumya Saurabh

**Affiliations:** Department of Chemistry, New York University, New York, New York 10003, United States; Department of Chemistry and Biochemistry, University of California, San Diego, La Jolla, California 92093-0358, United States

## Abstract

Aminoglycoside antibiotics bind RNA with high affinity through networks of amine and hydroxyl contacts, yet whether this multivalent binding can drive macroscopic RNA phase transitions has never been tested. Here we show that aminoglycosides are a class of small-molecule RNA condensers, and we take neomycin B (neoB), an FDA-approved member of the family, as a representative drug through which to dissect the mechanism. NeoB induces concentration-dependent phase separation of poly(A), poly(U), and total *E. coli* RNA, and kanamycin, apramycin, and gentamicin condense RNA as well. Condensate size and density are tunable by pH and ionic strength, which independently modulate neoB protonation and screening of interdroplet repulsion. NeoB forms more stable condensates than spermine despite spermine’s larger effective charge at physiological pH, whereas the amine-free polyol fucitol fails to condense RNA. Molecular dynamics simulations attribute neoB’s greater efficiency to additional hydrogen bonds donated by its hydroxyl groups. The chemical complexity that aminoglycosides evolved for RNA recognition thus also drives a macroscopic RNA phase transition that may contribute to bactericidal activity and cellular toxicity.

## Introduction

Macromolecular condensation organizes biomolecules into membraneless compartments that regulate metabolism, translation, and stress response across all domains of life.^1–3^ RNA molecules play active roles in condensate assembly: their polyanionic backbones provide numerous electrostatic contacts, while base pairing and stacking interactions tune condensate material properties.^4–6^ Polyamines such as spermine condense poly(U) RNA through complex coacervation, with phase behavior modulated by charge, crowding, and temperature-dependent reentrant transitions.^7,8^ Despite extensive biophysical characterization of protein-and ion-driven RNA condensates,^5,6^ no FDA-approved small-molecule drug has been shown to induce RNA phase separation in a quantitatively tunable manner.

Aminoglycosides are polycationic antibiotics that bind folded RNA with affinities in the nanomolar to micromolar range,^9–11^ yet their interactions with RNA have been characterized predominantly at the level of site-specific binding. Neomycin B (neoB) bears six primary amines and eight hydroxyl groups on a rigid aminocyclitol scaffold and binds the bacterial ribosomal A-site through a network of hydrogen bonds and electrostatic contacts.^12–15^ This recognition reflects the electrostatic complementarity between the polycationic antibiotic and the electronegative pocket of a folded RNA.^16,17^ Beyond site-specific recognition, aminoglycosides condense double-stranded DNA into toroids, rods, and spheres,^18^ and amphiphilic aminoglycoside derivatives are exploited as nucleic acid delivery agents. ^19^ In each case the nucleic acid collapses into a solid, condensed particle; whether an aminoglycoside can instead drive RNA into a liquid, tunable condensate had not been examined. Recent work has clarified that aminoglycosides promote mistranslation by stabilizing near-cognate tRNA accommodation,^13^ yet ribosomal binding alone does not account for their bactericidal potency or their characteristic ototoxicity and nephrotoxicity in mammalian cells.^20–22^ These off-target effects implicate mitochondrial dysfunction and RNA dysregulation, ^22,23^ but the molecular events connecting aminoglycoside–RNA binding to downstream phenotypes remain unclear.

Here we show that neomycin drives robust, concentration-dependent RNA phase separation across poly(A), poly(U), and total bacterial RNA extracts. We map this behavior across cation and RNA concentrations. Condensate material properties respond to pH and ionic strength through orthogonal tuning of protonation state and interdroplet repulsion. Comparison with spermine and fucitol establishes that cationic charge is necessary but not sufficient: neoB produces more stable condensates than spermine despite the latter carrying at least as much effective charge at physiological pH, pointing to a role for scaffold geometry and hydrogen bonding. These findings establish aminoglycoside–RNA condensation as a tunable, charge-driven phase transition. Its stability reflects the hydrogen-bonding and scaffold chemistry of the aminoglycoside rather than charge alone, connecting antibiotic structure to a physical transformation of RNA.

## Results and discussion

### Neomycin drives RNA phase separation

We mixed commercial neomycin sulfate (neoS; predominantly neomycin B, hereafter neoB; Figure S1) with poly(A) RNA in 10 mM Tris-HCl (pH 7.9, 50 mM NaCl, 30 ^◦^C) and imaged samples by differential interference contrast (DIC) microscopy. Phase separation was concentration dependent (Figure 1A). No condensates formed below approximately 50 µM neoS; above this threshold, microdroplets appeared and grew with neoS concentration. Condensation required both components: neoS alone produced no condensates, and droplets appeared only once poly(A) exceeded ∼0.1 µM. This stoichiometric co-dependence rules out precipitation by either component alone. Droplets fused on contact and relaxed into single spheres, confirming they were liquid (Figure S2B).^24^

**Figure 1:**
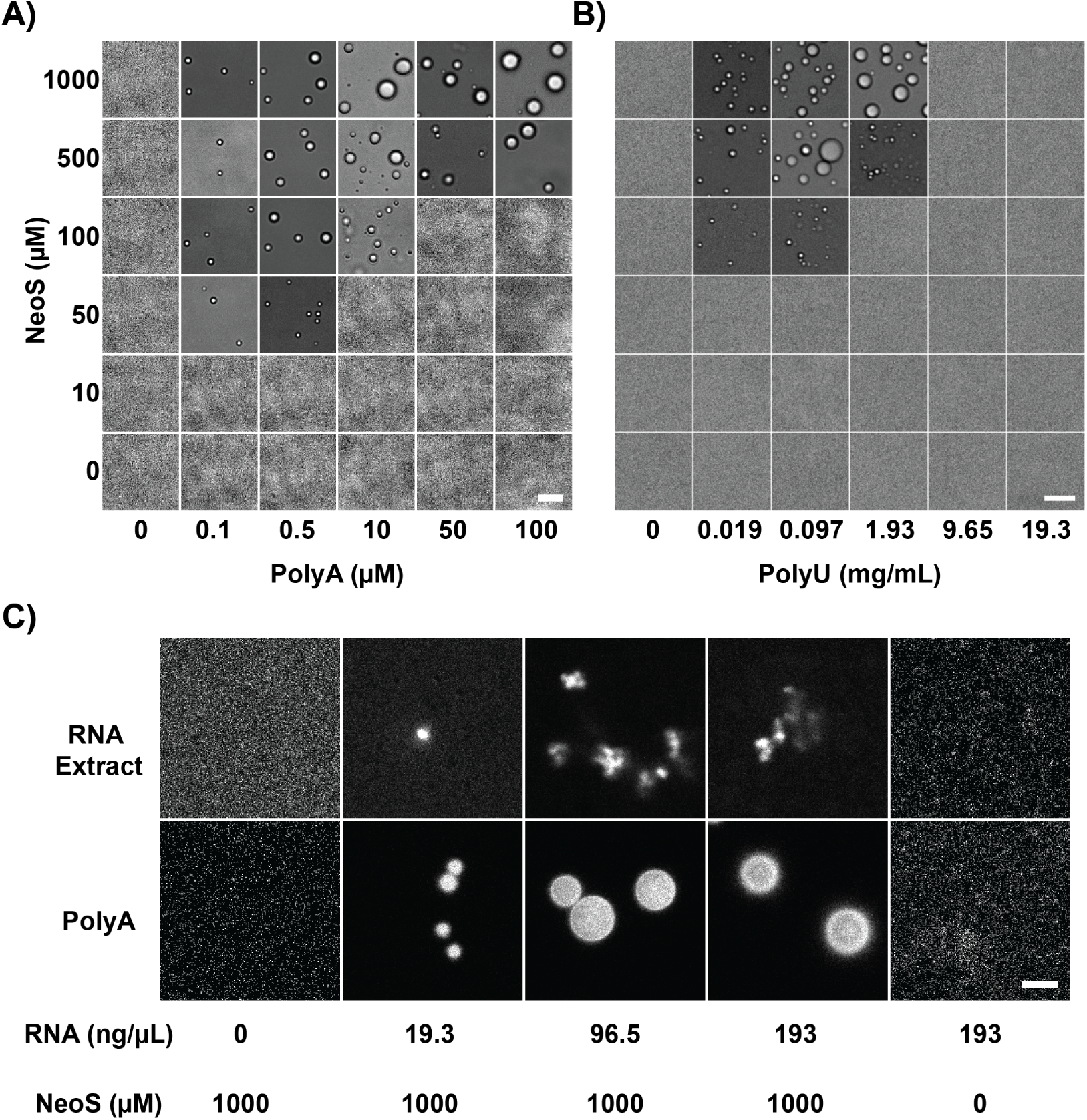
Neomycin sulfate induces phase separation across diverse RNAs. (A) DIC images of a two-dimensional phase diagram for poly(A) RNA (0–100 µM) and neomycin sulfate (neoS, 0–1000 µM) in 10 mM Tris-HCl pH 7.9, 50 mM NaCl, 30 ^◦^C. Scale bar, 5 µm. (B) DIC images of a two-dimensional phase diagram for poly(U) RNA (0–19.3 mg/mL) and neoS (0–1000 µM); a second, compositionally distinct RNA also condenses. Scale bar, 10 µm. (C) Phase separation of total *E. coli* RNA extract or poly(A) RNA at indicated concentrations with neoS at indicated concentrations. Samples were labeled with 10 µM Thioflavin T (ThT). Scale bar, 2 µm.

Condensation did not require a specific sequence. NeoS also drove phase separation of poly(U) RNA (Figure 1B), a homopolymer compositionally distinct from poly(A) but sharing the same polyanionic backbone. Because two sequence-distinct homopolymers both condense, condensation is driven by the shared polyanionic backbone rather than by sequence-specific recognition. Total RNA extracted from *E. coli* also condensed upon neoS addition, forming smaller and more irregular condensates than the homopolymers (Figure 1C). Condensation therefore generalizes from defined homopolymers to the heterogeneous RNA present in cells.

Condensation was also not unique to neomycin. Kanamycin, apramycin, and gentamicin each drove poly(A) phase separation in a concentration-dependent manner (Figure S2A), establishing RNA condensation as a general property of the aminoglycoside class rather than a peculiarity of a single antibiotic. The threshold concentration varied with the antibiotic, with droplets appearing by approximately 400 µM for gentamicin, 600 µM for apramycin, and 800 µM for kanamycin. Gentamicin also produced the largest droplets. This ordering follows the number of cationic amines each antibiotic carries—five each for gentamicin and apramycin against four for kanamycin, rising to six for neomycin B—rather than its hydroxyl content, which runs the opposite way: the hydroxyl-poor gentamicin condenses most readily while the hydroxyl-rich kanamycin condenses least. Across these scaffolds the cationic amine count, and thus the charge it confers, is therefore the primary determinant of condensation efficiency, and additional hydroxyl groups cannot offset fewer amines. Hydroxyl groups instead add a secondary enhancement only when charge is held constant, as the neoB–spermine comparison below establishes.

### Ionic strength and pH tune condensate size and density

If neoB drives condensation through its protonated amines, pH and ionic strength should act on distinct interactions. pH sets neoB’s protonation and thus the charge that drives its attraction to RNA, whereas added salt screens the surface charge that keeps droplets apart. We varied the two independently at fixed poly(A) and neoS, imaging by DIC and holographic microscopy, to distinguish the interactions that stabilize the condensed phase from those that oppose droplet coalescence (Figure 2).

**Figure 2:**
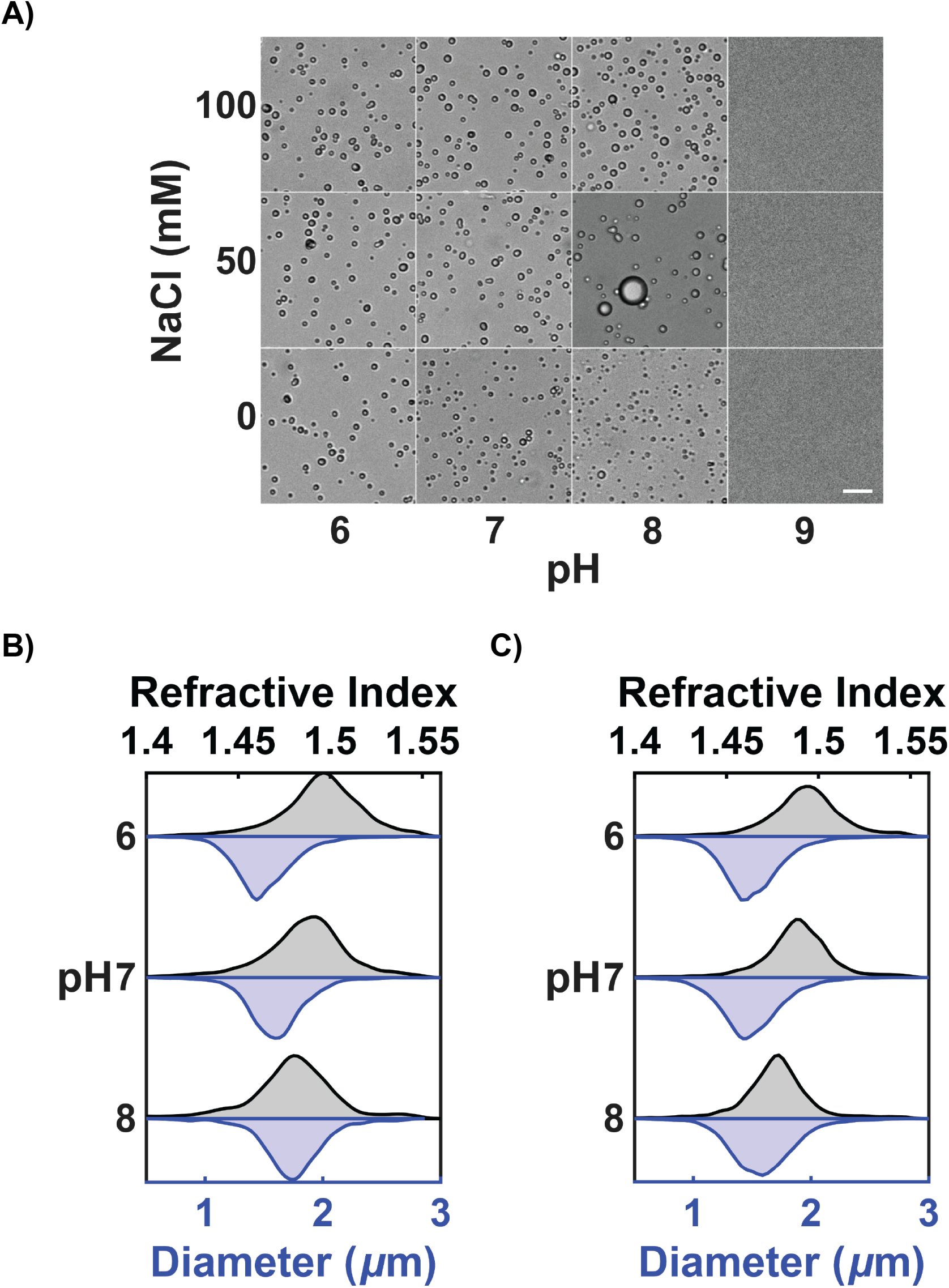
Ionic strength and pH tune condensate properties. (A) DIC images of poly(A) (1 µM) with neoS (200 µM) in 10 mM Tris-HCl at the indicated pH and NaCl concentration. Scale bar, 10 µm. (B and C) Size (blue) and refractive index (gray) distributions of condensates formed with 1 µM poly(A) and 200 µM neoS at the indicated pH with (B) 0 mM NaCl or (C) 100 mM NaCl.

DIC imaging across the pH and salt matrix showed that a protonated neoB was required for condensation and that salt set droplet size (Figure 2A). Condensates formed from pH 6 to 8 but were absent at pH 9, where neoB is largely deprotonated (Figure S3). At 0 mM NaCl, droplets were small and dispersed at every pH. Adding salt promoted coalescence into larger droplets, most conspicuously at pH 8 and 50 mM NaCl, the condition of lowest neoB charge. At 100 mM NaCl and acidic pH, droplets were numerous but small. Coalescence was thus greatest when the interdroplet barrier was low, and arrested when high neoB protonation raised it.

We used holographic microscopy to measure condensate size and interior density simultaneously and without fluorescent labeling.^25^ At 0 mM NaCl, diameters were narrow across pH 6–8, centered near 1 µm to 2 µm, and the refractive index rose as pH fell, indicating denser interiors at higher neoB protonation (Figure 2B). At 100 mM NaCl, diameters broadened at pH 7–8 while the refractive index stayed centered near 1.5 across pH 6–8, so interior density remained high even as the droplets coarsened (Figure 2C).

Two observations distinguish these condensates from a simple polyelectrolyte complex. First, the loss of condensation at pH 9 shows that the protonated amines are indispensable: no amount of salt restored condensation once neoB was deprotonated, so screening cannot substitute for the driving charge. Second, salt did not dissolve the condensates. Classical complex coacervates dissolve as added salt screens the attraction between polyions.^7^ Here condensates persisted from 0 to 100 mM NaCl, and salt instead tuned their coalescence. Salt therefore acts on the repulsion between droplets, not on the cohesion within them. Together, these give neoB two roles: its protonated amines cross-link the RNA backbone to hold the condensate together, tuned by pH, while its residual surface charge sets the barrier to droplet coalescence, tuned by salt.

### Neomycin condenses RNA at lower cation loading than spermine

Spermine is the canonical small-molecule condenser of RNA. ^7,8^ To test whether neoB’s advantage over it reflects more than its cationic charge, we compared the two across temperature, expressing each cation’s loading as the ionic strength it contributes (from the cation itself, not added NaCl) so that the comparison is charge-normalized (Figure 3).

**Figure 3:**
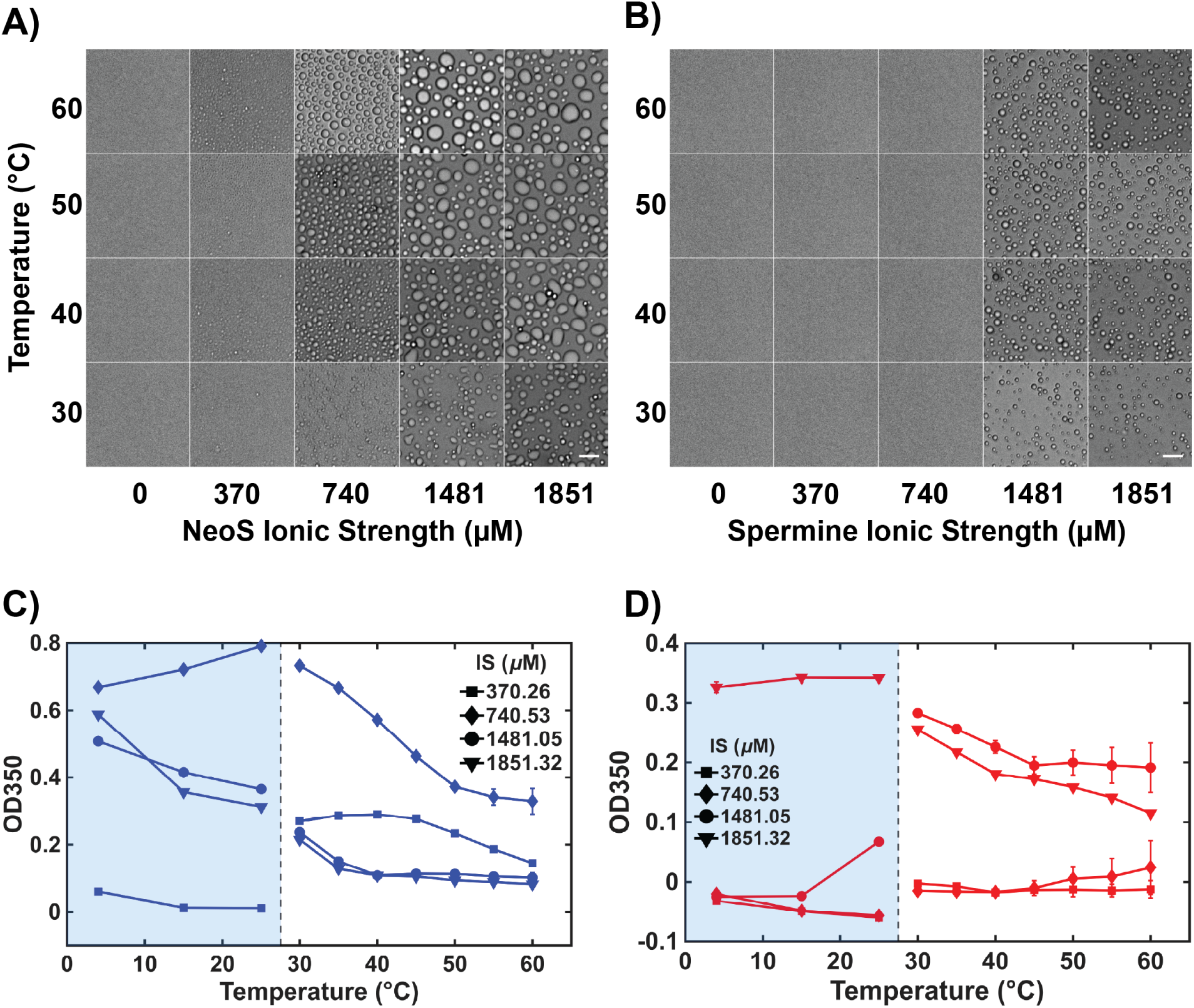
Poly(A) condensates persist over a broad temperature range. (A and B) Temperature-dependent morphology. Condensates were formed with 10 µM poly(A) and the indicated (A) neoS or (B) spermine ionic strength, then equilibrated at each temperature before imaging. Scale bars, 10 µm. (C and D) Turbidimetry (OD_350_) versus temperature for poly(A) condensates formed with (C) neoS or (D) spermine at the indicated ionic strengths. The measurements in the shaded region below 30 ^◦^C and those above it are separate experiments that were not cross-calibrated, and turbidity reports droplet number, size, and contrast rather than the dense-phase fraction; the two segments should not be read as a single continuous curve. Error bars are standard deviations of triplicate measurements.

NeoB condensed poly(A) at lower cation loading than spermine. NeoS formed condensates at an ionic strength of 740 µM, whereas spermine required roughly twice as much, 1481 µM, to condense poly(A) at all (Figure 3A,B). Above their thresholds both cations produced droplets that persisted to 60 ^◦^C, the highest temperature tested.

Turbidimetry quantified the difference (Figure 3C,D). At matched ionic strength, neoS condensates reached OD_350_ ∼0.8 versus below ∼0.34 for spermine, indicating denser or more numerous droplets. For neoS, turbidity peaked at an intermediate ionic strength (740 µM) and fell at higher loading, the reentrant signature of an electrostatically driven condensation.^8^ The direction of the temperature response depended on cation loading rather than following a single trend, consistent with the reentrant phase behavior reported for RNA–polycation coacervates along the temperature axis.^8^ We do not treat turbidity as an order parameter. OD_350_ reports droplet number, size, and refractive-index contrast, not the fraction of material in the dense phase, so it cannot define a critical temperature or separate increased phase separation from droplet coarsening. The data below 30 ^◦^C and above it were acquired as separate experiments that were not cross-calibrated, so the two segments should not be read as one continuous curve.

Two features point beyond simple charge neutralization. First, charge cannot account for the neoB–spermine gap: at the experimental pH 7.9 spermine carries the larger effective charge (∼+3.3 versus ∼+2.6 for neoB; Figure S3), yet neoB condenses at lower loading and to higher density. Second, the survival of the dense phase to 60 ^◦^C argues against a purely enthalpic ion-pairing mechanism. Associative phase separation of polyelectrolytes is driven largely by the entropy released when condensed counterions and ordered water are freed on complex formation, a contribution that grows with temperature.^7,8^ This solvent-and counterion-entropy effect is distinct from the ligand conformational entropy considered below. The thermal robustness we observe is consistent with such an entropically stabilized condensate. Structural features beyond the amines must therefore underlie this difference. To identify them, we compared neoB–poly(A) and spermine–poly(A) interaction ensembles by molecular dynamics simulation, focusing on direct hydrogen bonding, ligand-mediated bridging, and representative bound conformations.

### Hydroxyl groups and a rigid scaffold favor neomycin over spermine

To separate charge from hydrogen bonding, we compared two chemically minimal probes: spermine, which bears amines but no hydroxyl groups, and fucitol, a hexitol with hydroxyl groups but no amines. Spermine (0–250 mM) drove poly(A) phase separation (Figure 4A), whereas fucitol produced no condensates at any condition tested (0–1000 mM fucitol, 0–100 µM poly(A); data not shown). Protonatable amines are necessary for condensation; hydroxyl groups alone are not.

**Figure 4:**
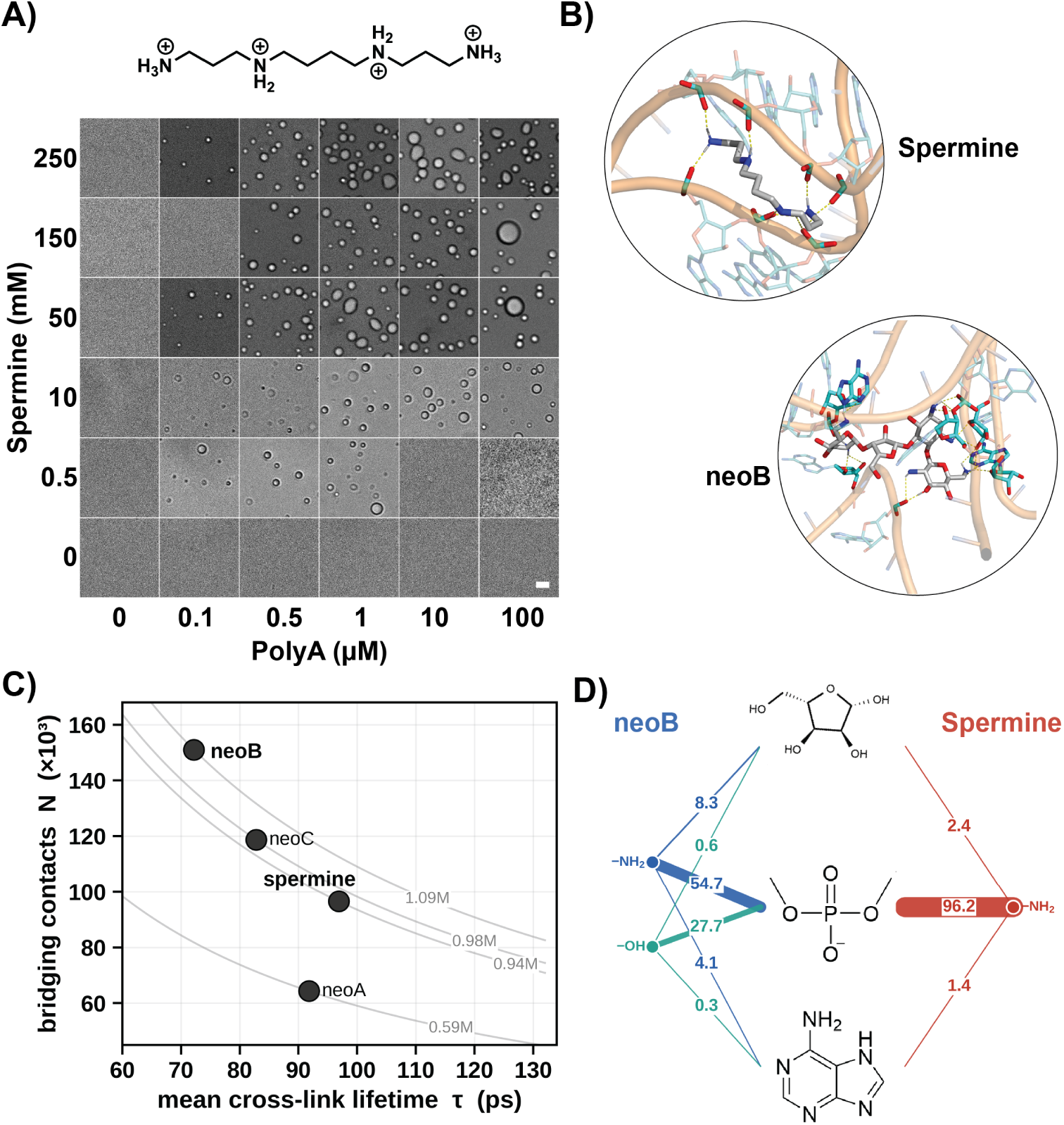
Amines drive poly(A) condensation, and hydroxyl contacts distinguish neoB from spermine. (A) DIC images of a two-dimensional phase diagram for poly(A) (0–100 µM) with spermine (0–250 mM); spermine drives phase separation in a concentration-dependent manner. Scale bar, 5 µm. (B) Representative molecular dynamics snapshots of spermine (top) and neoB (bottom) bridging poly(A); dashed lines indicate direct hydrogen bonds. (C) Ligand-mediated bridging of poly(A) by each protonated cation: number of bridging contacts *N* against mean cross-link lifetime *τ*, with gray curves marking iso-occupancy (*Nτ* = constant). NeoB forms many short-lived cross-links and spermine fewer, longer-lived ones at similar total occupancy. (D) Hydrogen-bond map from ligand donor groups to poly(A) acceptor moieties for neoB (left) and spermine (right). Line thickness is proportional to the time-weighted hydrogen-bond occupancy (%; value labelled on each edge); blue, neoB amine (−NH_2_) donor; teal, neoB hydroxyl (−OH) donor; red, spermine amine donor. Central structures are the poly(A) acceptor moieties (phosphate, ribose, adenine). Both cations bond predominantly to the phosphate backbone, but only neoB also donates hydrogen bonds from its hydroxyl groups.

This requirement is echoed by the loss of condensation when neoB is deprotonated at pH 9. It does not, however, explain why neoB condenses poly(A) at lower loading than spermine, whose effective charge is comparable (Figure 3). We therefore asked whether neoB’s hydroxyl groups act together with its amines, adding hydrogen bonds that spermine cannot make.

Purified neoB reproduced poly(A) condensation (Figure S4), confirming that condensation is intrinsic to neoB and not an artifact of the sulfate counterion. Purified neoB, however, phase separated only at a higher concentration than neoS, with a correspondingly higher saturation concentration across the poly(A) range tested (Figure S4B). The lower saturation concentration of neoS most likely reflects two complementary contributions that purified neoB lacks. Its divalent, kosmotropic sulfate counterion, in place of the trifluoroacetate of the purified salt, favors phase separation at lower cation loading; and its neoA and trace neoC congeners (Figure S1) themselves bridge poly(A) in the molecular dynamics simulations below. The two effects act in the same direction, and we therefore included neoA and neoC, alongside neoB and spermine, in those simulations.

Molecular dynamics simulations of neoB– and spermine–poly(A) ensembles resolve the role of the hydroxyl groups (Figure 4B–D; Figures S5 and S6). In the protonated state, both cations bind mainly the phosphate backbone rather than the bases: over half of neoB’s direct hydrogen bonds are made to phosphate oxygens, consistent with the backbone-driven condensation seen for the homopolymers (Figure 1). Only neoB also donates hydrogen bonds from its hydroxyl groups, which account for about a third of its hydrogen-bond occupancy and have no counterpart in spermine (Figure 4D). NeoB also engages the ribose and adenine moieties far more than spermine, with hydrogen bonds to these base and sugar sites accounting for 14.8 % of its occupancy versus 3.8 % for spermine. At matched effective charge, the two cations reach similar total bridging occupancy by opposite routes: neoB forms many short-lived cross-links and spermine fewer, longer-lived ones, so neoB bridges through contact number rather than contact strength (Figure 4C). Cross-linking also requires the cationic groups to adopt an extended, distributed geometry (Figure S5): a flexible polyamine samples this bridging-competent arrangement only transiently and at a conformational entropy cost, whereas the rigid aminoglycoside scaffold presents its amines pre-organized in that geometry. NeoB and spermine differ in several properties at once, including amine number, primary versus secondary amine character, hydroxyl content, and scaffold rigidity, so the present data bound rather than isolate the contribution of any single feature to neoB’s advantage.

Together the phase behavior, turbidimetry, holographic imaging, and simulations support a layered model: amine–phosphate electrostatics and the entropy of counterion release provide the driving force, hydroxyl hydrogen bonds to the RNA backbone add binding that spermine lacks, and the residual surface charge sets the barrier to droplet coalescence. The model predicts that changing amine count or hydroxyl geometry across the aminoglycoside family will shift the saturation concentration.

### Discussion and outlook

The data presented here establish that neomycin B drives phase separation of polyanionic RNA. The phase boundaries are concentration dependent. The condensates are liquid-like, undergoing fusion and coarsening over time. Their size, density, and coalescence respond to pH, ionic strength, and the identity of the condensing agent. Comparison with spermine and the negative control with fucitol locates the driving force in the amine groups, while neoB condenses more readily because of additional hydrogen bonds from its hydroxyl groups and the rigidity of its scaffold. This interpretation is consistent with both simulation and structural precedent. In the MD trajectories, neoB sampled more bridge-rich geometries than spermine. Aminoglycosides are best known for specific recognition of a single folded RNA: at the ribosomal decoding site, neomycin B clamps one duplex through the conserved A1408/A1492/A1493 triad (PDB 2ET4).^26^ Yet the same scaffold can bridge two separate RNA chains when geometry permits, spanning the two hairpins of the HIV-1 dimerization-initiation-site kissing loop (PDB 2FCY)^27^ and contacting multiple RNA chains in a dedicated neomycin–RNA crystal (PDB 2A04).^28^ These structures show that the neomycin B scaffold can clamp two RNA chains at once. In our condensates the same bridging recurs many times along the polyanionic backbone and without sequence specificity, so that repeated transient cross-links drive a macroscopic phase transition.

Total *E. coli* RNA extract condenses under neoB at concentrations comparable to poly(A). Inside a bacterial cell, rRNA accounts for approximately 80% of total RNA and reaches local concentrations of hundreds of micromolar. ^21,29^ Intracellular aminoglycoside concentrations during clinical treatment reach tens to hundreds of micromolar through proton motive force driven uptake.^21,30^ The phase boundaries we map in vitro fall within physiologically accessible conditions. RNA condensation in drug-treated bacteria would concentrate and immobilize the translational machinery, offering a parallel mechanism of bactericidal action alongside ribosomal mistranslation.^20,21^

Aminoglycosides preferentially accumulate in cochlear hair cells and renal proximal tubular cells. The intracellular events leading to irreversible damage in these cells have not been explained by mammalian ribosomal RNA targeting alone.^22,23^ Both cell types are among the most translationally active in the body and sustain high levels of rRNA and mRNA. We propose that RNA condensation by accumulated aminoglycosides may sequester essential transcripts, reduce ribosome accessibility, or alter the properties of endogenous RNA granules such as stress granules and P-bodies, contributing to the irreversible cellular damage that defines aminoglycoside toxicity. This hypothesis offers a testable framework linking aminoglycoside–RNA condensation to cellular toxicity.

In bacteria, condensation has so far been studied mainly as a protein-driven response to physiological cues.^3,31^ The present work extends that scope: a small-molecule antibiotic, rather than an intrinsically disordered protein, drives condensation, adding chemical perturbation by exogenous drugs to bacterial condensate biology.

The generality across kanamycin, apramycin, and gentamicin (Figure S2A) already indicates that condensation is a class property whose efficiency tracks antibiotic structure. Key questions now follow: whether these condensates form in living bacteria at therapeutic doses and affect translation, whether structurally distinct aminoglycosides condense RNA in proportion to their amine and hydroxyl content, and whether condensation can be decoupled from ribosomal binding by scaffold modification.

## Experimental section

### Materials

Poly(A) potassium salt (average MW ∼193,000, ∼556 nt) and poly(U) potassium salt were purchased from Sigma-Aldrich (P9403 and P9528, respectively). Neomycin sulfate (USP grade) was from Thermo Fisher Scientific (Gibco, cat. no. 21810-031). Spermine tetrahydrochloride was from Sigma-Aldrich (S1141). L-Fucitol was from Santa Cruz Biotechnology. Thioflavin T (ThT) was from Sigma-Aldrich (T3516). All buffers were prepared using nuclease-free water (Invitrogen). Total RNA was extracted from mid-log phase *E. coli* (serotype O157:H7) using TRI Reagent ™(Zymo Research Corp.).

### Neomycin composition and purification

The composition of commercial neomycin sulfate was determined by liquid chromatography–mass spectrometry. Chromatographic separation was performed on a Thermo Scientific TSQ Fortis triple-quadrupole liquid chromatography–mass spectrometry (LC-MS) system equipped with an autosampler maintained at 10 °C. Separation was carried out on a Varian Microsorb-MV 100-5 C18 column (250 × 4.6 mm, 5 µm particle size) at ambient temperature, under isocratic conditions using a mobile phase of water/methanol (40:60, v/v) containing 1.7 mL/L heptafluorobutyric acid (HFBA) as an ion-pairing reagent, delivered at a flow rate of 0.4 mL/min over a total run time of 45 min. The injection volume was 10 µL. Samples were dissolved in water containing 1.7 mL/L HFBA to prepare a stock solution at 0.5 mg/mL. The solutions were filtered through a 0.22 µm membrane filter prior to injection. Due to its comparatively low ionization response, neomycin A was analyzed at a higher concentration of 5 mg/mL to enable reliable detection. Mass spectrometric detection was performed in positive electrospray ionization (ESI^+^) mode. Full-scan mass spectra were acquired over *m/z* 100–1000, and selected ion monitoring (SIM) was used to extract the protonated molecular ions of neomycin B/C ([M+H]^+^, *m/z* 615) and neomycin A ([M+H]^+^, *m/z* 323) for identification and quantitation. Data acquisition and processing were performed using Thermo Scientific FreeStyle software (version 1.9 SP3). An Ambeed neomycin B/C reference was used as an analytical standard to resolve neomycin B from its co-eluting epimer neomycin C; by this analysis, neoS was 99.5 % neomycin B (neoB), 0.5 % neomycin A (neoA), and only trace neomycin C (Figure S1). Neomycin B was purified from the commercially available neomycin sulfate by Boc protection of the amines of the neomycin isomers, separation by silica gel column chromatography, and Boc removal^32^ of the isolated neomycin B to form the TFA salt. The identity and purity of the product were assessed by ^1^H NMR (Figure S4C).

### Phase separation assays

Stock solutions of poly(A) were prepared in 10 mM Tris-HCl pH 7.9 with 50 mM NaCl (unless otherwise stated) at 100 µM, aliquoted, and stored at −80 ^◦^C. For phase diagram experiments, RNA and cation solutions were mixed at the indicated concentrations in PCR tubes and incubated for 1 hour at 30 ^◦^C. Samples were transferred to 1% Tween-20-treated glass-bottom 384-well plates (Cellvis) for imaging. To treat well plates, the wells were first flushed with N_2_ gas to remove dust and debris, then coated with 1 % Tween-20 and incubated at 37 ^◦^C for 1 h. After incubation, the wells were washed five times with water at twice the volume of the Tween-20 solution. Finally, the wells were dried with N_2_ gas and used within the same day. For pH and ionic strength experiments, buffers were adjusted with NaOH/HCl and supplemented with NaCl as indicated. Temperature experiments involved incubation at 30 ^◦^C for 1 hour followed by 30 minutes at the target temperature before imaging above 30 ^◦^C or incubated for 30 min at the indicated temperature below 30 ^◦^C.

### Turbidimetry

Absorbance at 350 nm was measured using a plate reader (Cytation 5). Samples contained 10 µM poly(A) and the indicated cation concentrations in 10 mM Tris-HCl pH 7.9 with 10 mM NaCl. Temperature experiments involved incubation in PCR tubes at 30 ^◦^C for 1 hour followed by 30 minutes at the target temperature before imaging above 30 ^◦^C or incubated for 30 min at the indicated temperature below 30 ^◦^C before imaging. Measurements were blanked to a buffer-only control and performed in triplicate.

### Brightfield, DIC, and confocal microscopy

Brightfield images were collected using a multi-well plate imager (Cytation 5) equipped with a 60× air objective pre-heated to 30 ^◦^C. The use of a plate reader enabled rapid phase space imaging and phase boundary determination. DIC images were collected using an inverted microscope (Nikon Ti2) equipped with DIC optics, an oil immersion objective (Nikon PlanApo, 100×, 1.45 NA), and a sCMOS camera (Photometrics Prime 95B) with a system magnification of 0.11 µm*/*pixel. Confocal imaging of 10 µM ThT-labeled poly(A) or RNA extract condensates was performed using a confocal laser scanning microscope (Abberior). Images were acquired approximately 15 min after adding the samples to the well plate to allow condensates to settle.

### Holographic microscopy

Holographic imaging was performed using a commercial instrument (Spheryx xSight) as previously described.^33,34^ Briefly, condensate samples were transferred to microfluidic channels where pressure-driven flow transported the condensates. The dispersed particles were imaged with a 450 nm laser to produce holograms. The holograms were fit by the instrument using Lorenz-Mie theory to extract both refractive index and diameter of each particle.^34–36^

### Molecular dynamics simulations

#### System construction

Each system comprised a single poly(A) strand of 556 adenosine residues with ten ligand molecules, approximating the experimental poly(A)/ligand stoichiometry. To separate the contribution of cationic charge from scaffold chemistry, and because neoS is a mixture of neomycin congeners (Figure S1), we simulated spermine and neomycins A, B, and C, each in a neutral and a fully protonated state (spermine and neomycin A, 0 and +4; neomycin B and neomycin C, 0 and +6). RNA was described with the Amber OL3 (*χ*OL3) force field,^37–39^ and the ligands with the General Amber Force Field 2 (GAFF2),^40^ with atomic partial charges from AM1-BCC.^41^ Systems were solvated with TIP3P water^42^ in a truncated octahedral box and neutralized with Na^+^ counterions using the Joung–Cheatham monovalent ion parameters under Ewald electrostatics;^43^ no additional salt was added.

#### Equilibration and production

Simulations were performed with Amber 24 using the GPU-accelerated PMEMD engine, pmemd.cuda. Long-range electrostatics were evaluated by particle-mesh Ewald,^44^ bonds to hydrogen were constrained with SHAKE (ntc=2, ntf=2), and a 1 fs time step was used. Systems were energy-minimized in stages, heated from 10 K to 298.15 K, under restrained NVT conditions, equilibrated under restrained NPT conditions, and then relaxed at fixed volume before an unrestrained production run of 50 ns at 298.15 K. Production simulations were performed in the NPT ensemble (ntb=2, ntp=1) with Langevin temperature regulation (ntt=3, gamma_ln=2.0 ps^−1^) and pressure coupling to 1 atm (taup=10 ps). A 10.0 Å real-space nonbonded cutoff was used, and all-atom coordinates were saved every 10 ps, yielding 5000 frames per system.

#### Interaction analysis

Production trajectories were processed with AmberTools cpptraj.^45^ Trajectory segments were concatenated, and periodic boundary artifacts were corrected using autoimage before subsequent analysis. Ligand–poly(A) interactions were then classified on a per-frame basis using custom Python scripts that used the Amber topology and processed trajectory coordinates; NumPy and SciPy were used for geometric calculations. Contacts were classified into: overall association (any heavy-atom pair within 4.5 Å); phosphate–amine backbone contact (amine N within 4.0 Å of a phosphate oxygen); direct hydrogen bonds (donor–acceptor ≤3.5 Å, angle ≥ 120^◦^), resolved by donor type (amine or hydroxyl) and acceptor moiety (phosphate, ribose, adenine); amine–*π* contact (amine N within 6.0 Å of an adenine ring centroid); water-mediated contact; and ligand-mediated bridging (one ligand simultaneously contacting two or more distinct poly(A) residues). For each interaction class *k*, discrete contact events *i* were identified as contiguous trajectory frames satisfying the corresponding geometric criterion. The residence time *τ_i_* of each event was calculated from the number of consecutive frames in that event and the 10 ps frame spacing. The time-weighted occupancy was defined as

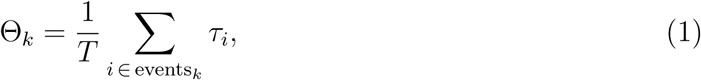

where *T* = 50 ns is the total analyzed production time for each system. Thus, Θ*_k_* represents the fraction of the production trajectory during which interaction class *k* was present. The mean residence time for interaction class *k* was calculated as

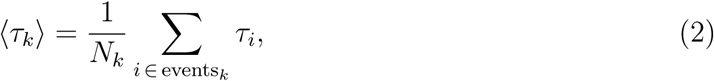

where *N_k_*is the number of discrete contact events.

#### Bridging conformational analysis

Bridging ligands were classified by the number of distinct poly(A) residues contacted (non-bridge, 0–1; bridge, ≥2; strong bridge, ≥4). For each ligand and frame the pairwise distances between its cationic nitrogen atoms were computed; mean distances in the bridging states were compared to the non-bridge state (distance-shift analysis), and the standardized distance vectors were projected by principal-component analysis to relate ligand conformation to bridging state (Figure S5).

## Acknowledgement

The authors thank members of Saurabh research group for helpful discussions. This study was supported by the National Institutes of Health through award 1R35GM157103 to S.S. and R35GM139407 to Y.T., the Margaret Strauss Kramer fellowship to J.vH, NYU Dean Undergraduate research funds to CC and KWL. This work was supported in part through the NYU IT High Performance Computing resources, services, and staff expertise. The Spheryx xSight holographic characterization instrument used in this study was acquired with support from the NSF under Award No. DMR-1420073.

## Supporting Information Available

Supplementary Figures S1–S6.

## Supporting information

**Figure S1:**
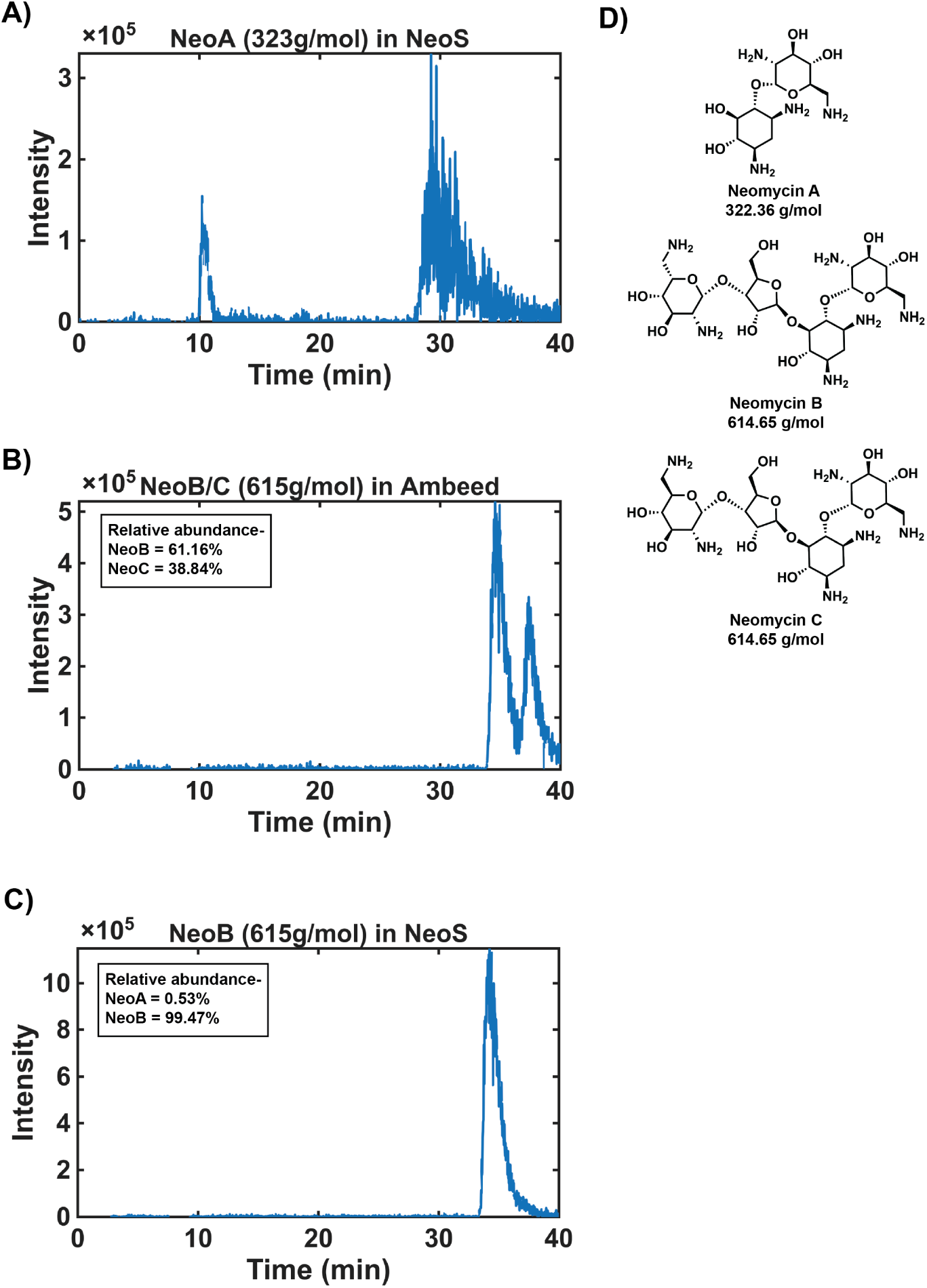
Liquid chromatography–mass spectrometry composition of the neomycin sources. (A) Extracted-ion chromatogram for neomycin A (neoA, 323 g/mol) in the neomycin sulfate (neoS) used in this study. (B) Neomycin B/C (615 g/mol) in an Ambeed neomycin standard, quantified as 61.2% neoB and 38.8% neoC; this B/C mixture serves as an analytical standard that resolves neomycin B from its co-eluting epimer neomycin C. (C) The 615 g/mol species in neoS, quantified as 99.5% neoB and 0.5% neoC; against the retention established in (B), this peak is essentially all neoB with only trace neoC. (D) Structures and molecular weights of neomycins A, B, and C.

**Figure S2:**
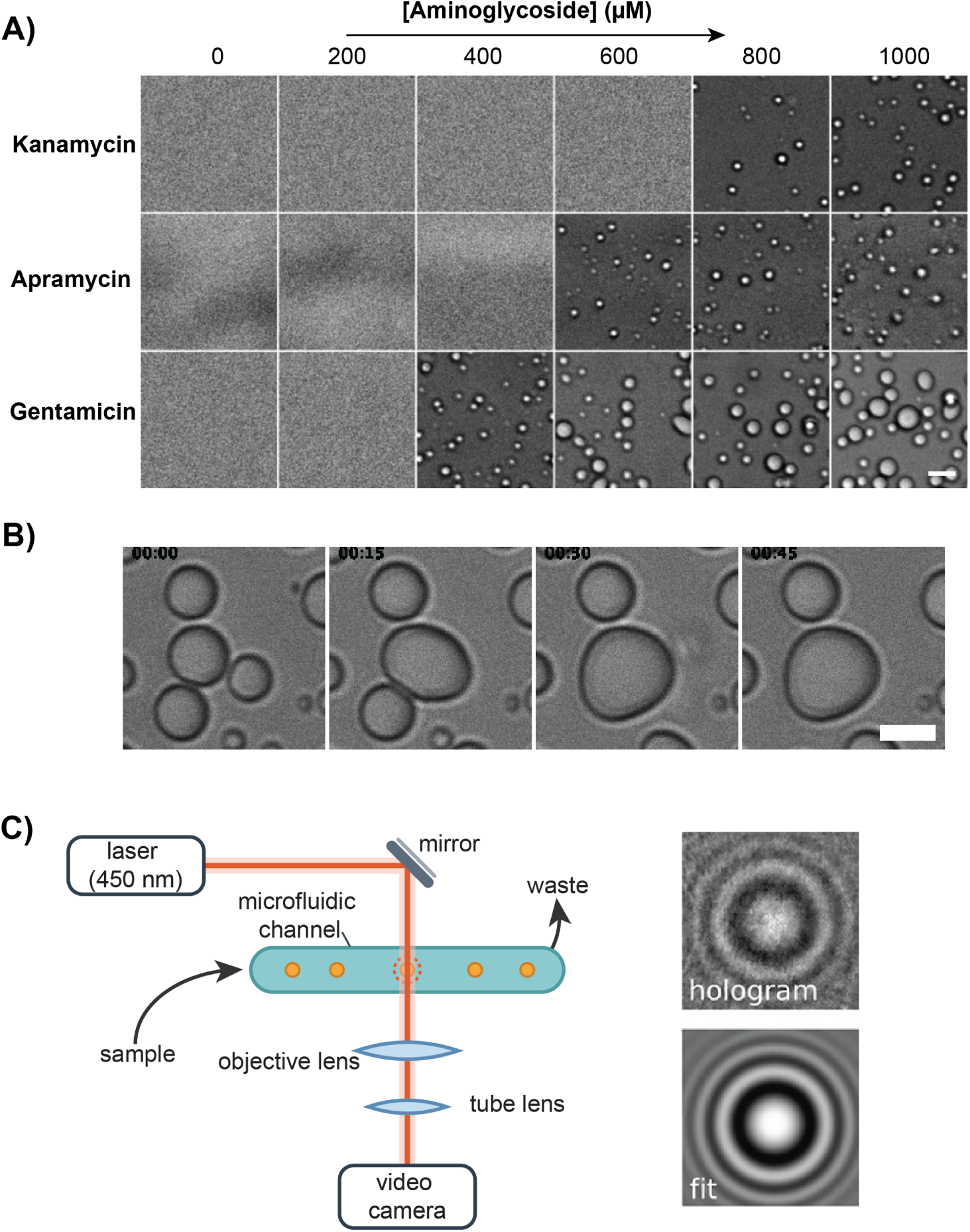
RNA condensation generalizes across the aminoglycoside class, and neomycin–poly(A) condensates are liquid-like. (A) DIC images of poly(A) (1 µM) mixed with kanamycin, apramycin, or gentamicin at the indicated concentrations (0–1000 µM) in 10 mM Tris-HCl pH 7.9, 50 mM NaCl, 30 ^◦^C. All three aminoglycosides drive concentration-dependent poly(A) phase separation; condensates appear at progressively lower antibiotic concentration for kanamycin, apramycin, and gentamicin. Scale bar, 5 µm. (B) Time-lapse DIC images showing two poly(A)–neomycin condensates fusing and relaxing into a single spherical droplet, confirming liquid-like behavior. Scale bar, 2 µm. (C) Schematic of holographic particle characterization: particles carried by pressure-driven flow through a microfluidic channel are illuminated by a laser, and each recorded hologram is fit to a Lorenz–Mie light-scattering model to extract particle diameter and refractive index.

**Figure S3:**
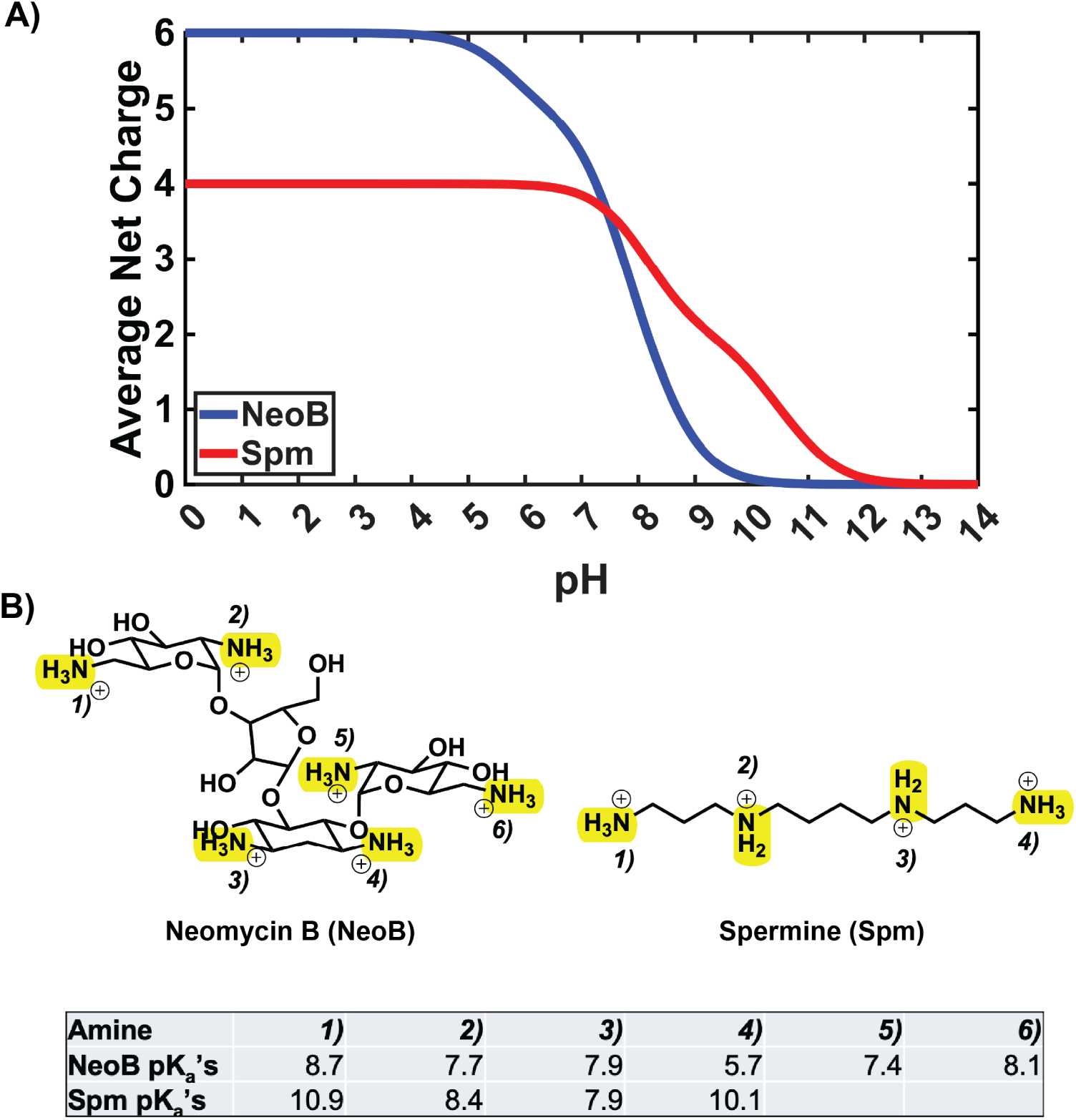
Protonation states of neoB and spermine as a function of pH. (A) Average net charge of neoB (blue) and spermine (Spm, red) versus pH, computed from the Henderson–Hasselbalch equation and literature p*K*_a_ values.^46^ The curves cross near pH 7.5, so at the experimental pH 7.9 spermine carries the larger effective charge (∼+3.3 versus ∼+2.6). (B) Structures of neoB and spermine with protonatable amines highlighted and numbered. (C) Literature p*K*_a_ value for each amine.

**Figure S4:**
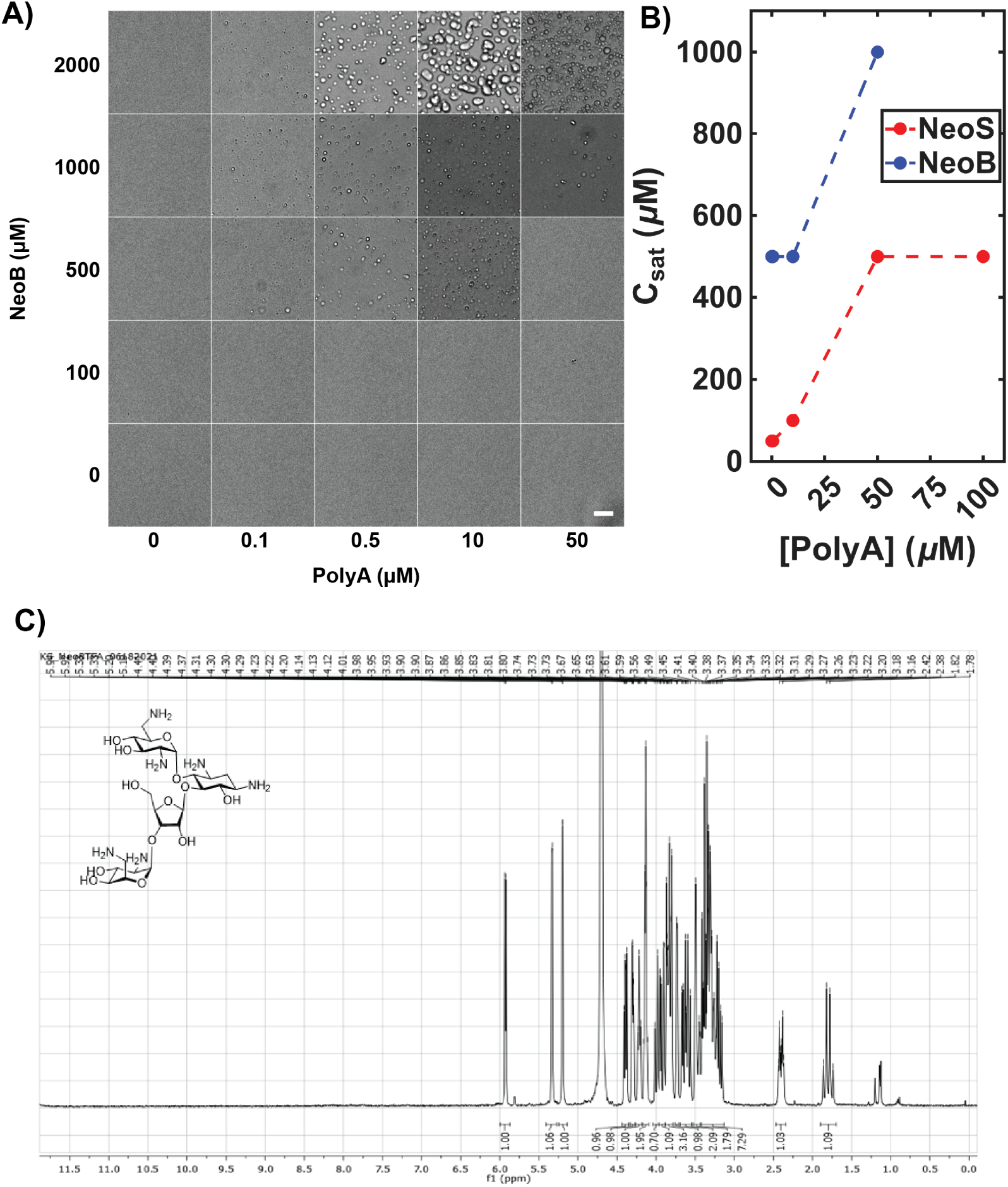
Purified neomycin B recapitulates poly(A) phase separation. (A) DIC images of a two-dimensional phase diagram for poly(A) (0–50 µM) and purified neoB (0–2000 µM) in 10 mM Tris-HCl pH 7.9, 50 mM NaCl, 30 ^◦^C. Scale bar, 10 µm. (B) Saturation concentration (*C*_sat_) versus poly(A) concentration for commercial neomycin sulfate (neoS, red) and purified neoB (blue). (C) ^1^H NMR (300 MHz, D_2_O) spectrum of the purified neomycin B (trifluoroacetate salt), confirming its identity and purity.

**Figure S5:**
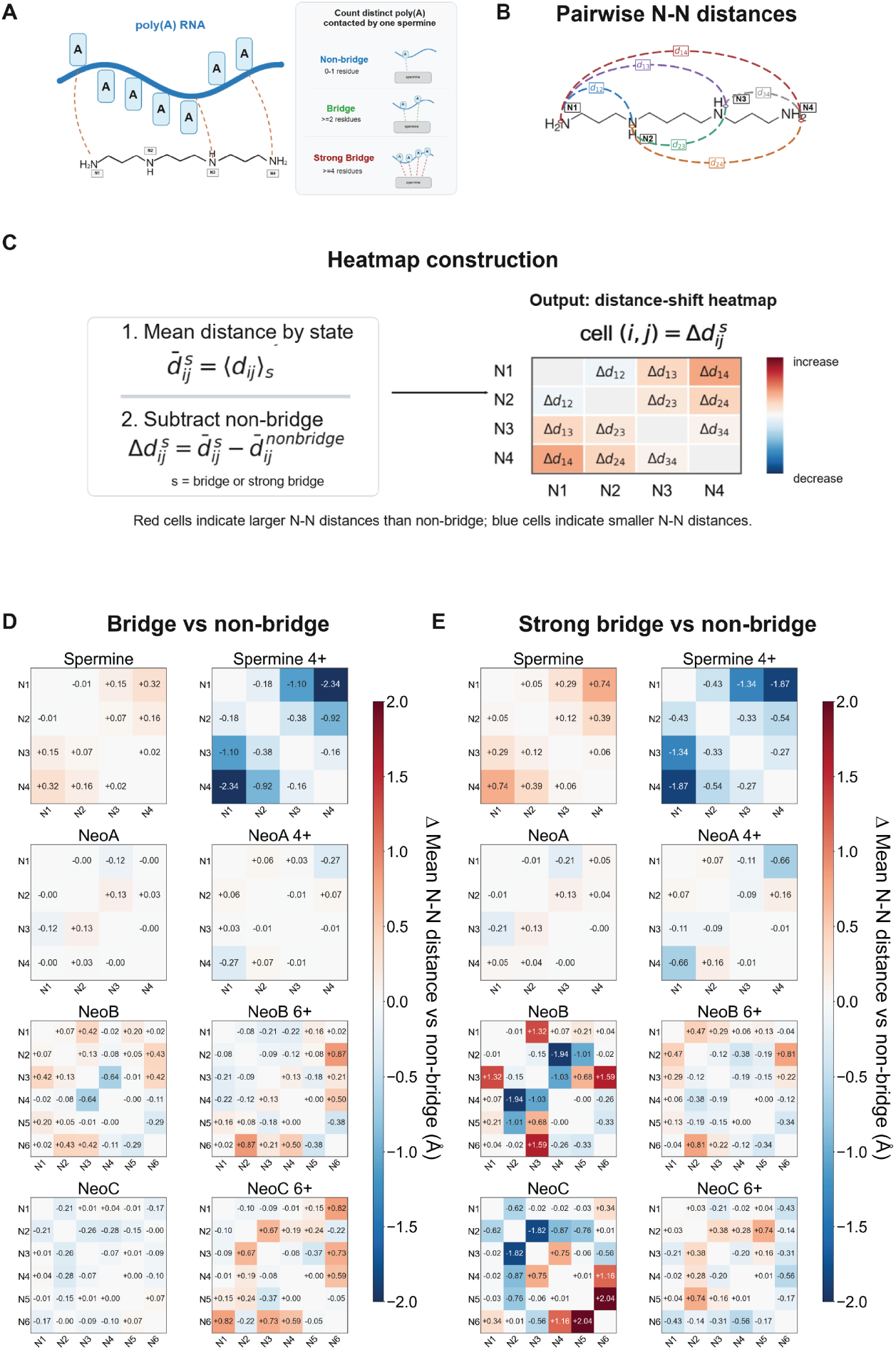
Bridging is accompanied by an extended cationic geometry: pairwise N–N distance analysis. (A) Bridging states defined by the number of distinct poly(A) residues contacted by a single ligand (non-bridge, 0–1; bridge, ≥2; strong bridge, ≥4). (B) The pairwise nitrogen–nitrogen (N–N) distances used to describe ligand conformation (illustrated for spermine). (C) Construction of the distance-shift heatmap: for each bridging state, the mean N–N distance minus its non-bridge value 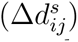; red denotes larger (extended) and blue smaller N–N distances. (D and E) Distance-shift heatmaps for the (D) bridge and (E) strong-bridge states, for spermine, neoA, neoB, and neoC in the neutral and fully protonated forms. Red cells mark N–N distances that increase upon bridging; the effect is largest for the strongly bridging, protonated ligands.

**Figure S6:**
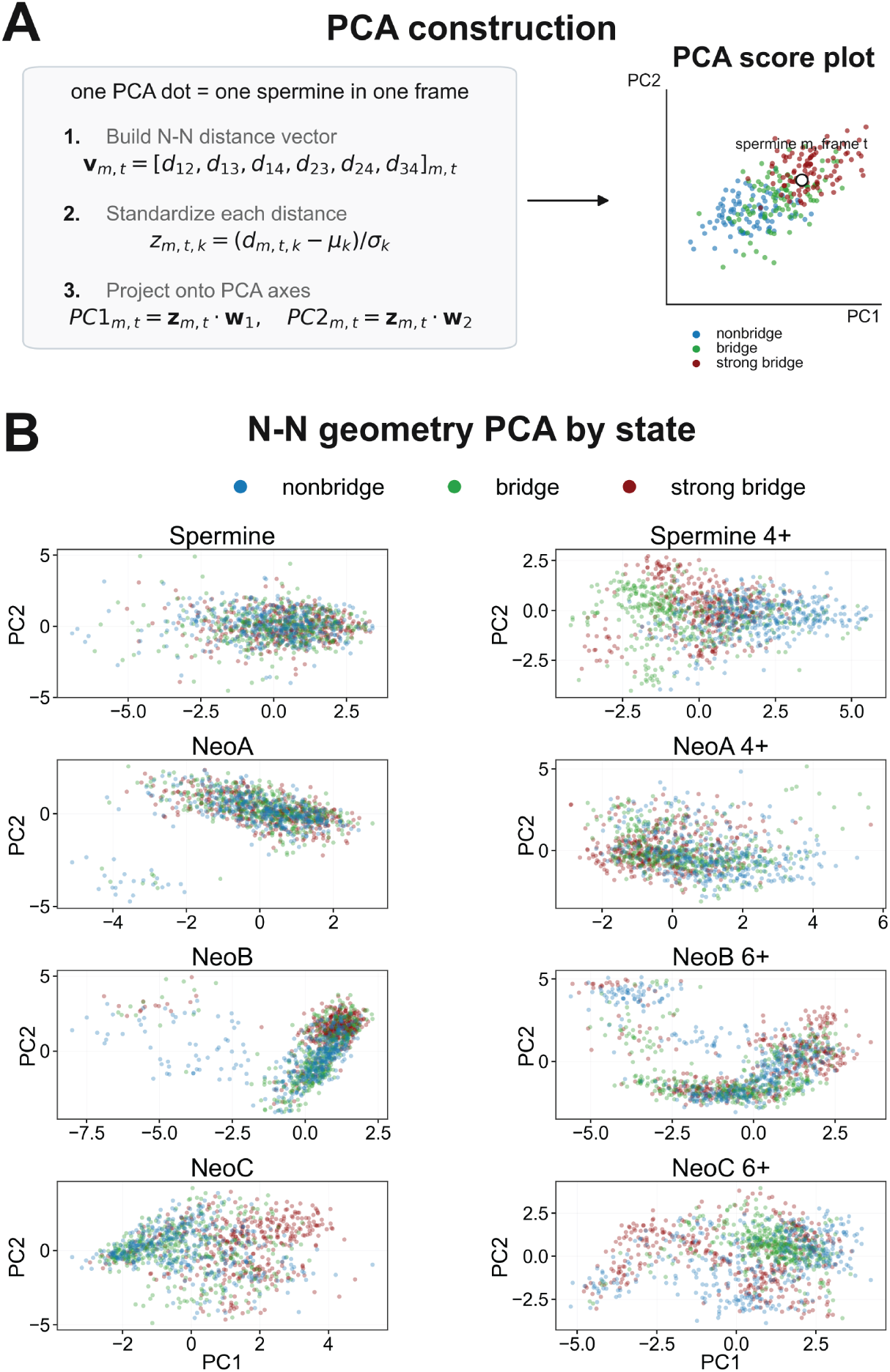
Principal-component analysis of ligand conformation separates bridging states. (Top) Construction of the analysis: for each ligand and frame the standardized N–N distance vector is projected onto its first two principal components; each point is one ligand in one frame, colored by bridging state (non-bridge, blue; bridge, green; strong bridge, red). (Bottom) PCA score plots for spermine, neoA, neoB, and neoC in the neutral and fully protonated forms. Strong-bridging frames occupy a distinct, more extended region of conformational space, most clearly for spermine^4+^, neoB, and neoC.

## TOC Graphic

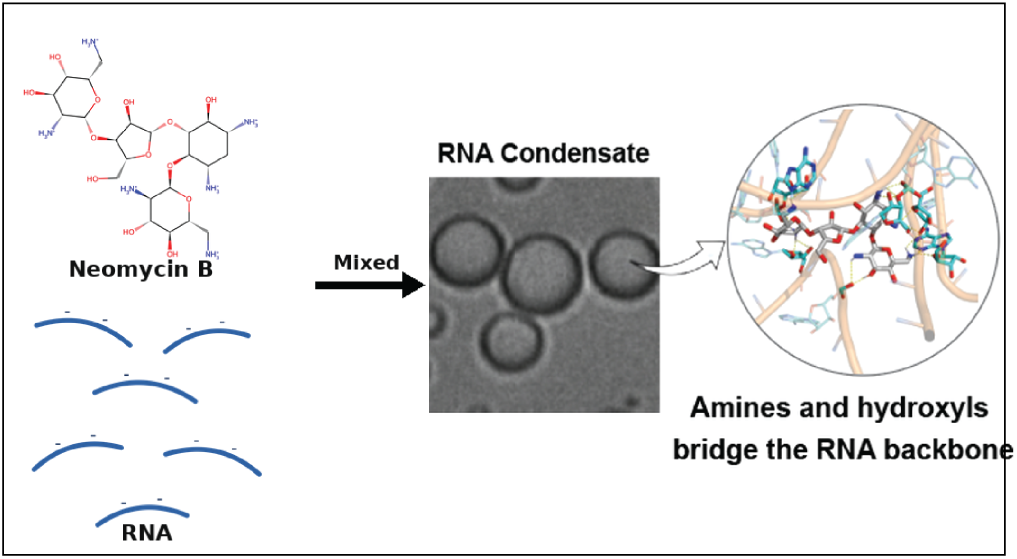

## References

(1) Banani, S. F.; Lee, H. O.; Hyman, A. A.; Rosen, M. K. Biomolecular condensates: organizers of cellular biochemistry. Nat. Rev. Mol. Cell Biol. 2017, 18, 285–298.

(2) Mittag, T.; Pappu, R. V. A conceptual framework for understanding phase separation and addressing open questions and challenges. Mol. Cell 2022, 82, 2201–2214.

(3) Saurabh, S.; Chong, T. N.; Bayas, C.; Dahlberg, P. D.; Cartwright, H. N.; Moerner, W. E.; Shapiro, L. ATP-responsive biomolecular condensates tune bacterial kinase signaling. Sci. Adv. 2022, 8, eabm6570.

(4) Lin, Y.; Fang, X. Phase separation in RNA biology. J. Genet. Genomics 2021, 48, 872–880.

(5) Roden, C.; Gladfelter, A. S. RNA contributions to the form and function of biomolecular condensates. Nat. Rev. Mol. Cell Biol. 2021, 22, 403–420.

(6) Jalihal, A. P.; Pitchiaya, S.; Xiao, L.; Bawa, P.; Jiang, X.; Bedi, K.; Parolia, A.; Cieslik, M.; Ljungman, M.; Chinnaiyan, A. M.; Walter, N. G. Multivalent proteins rapidly and reversibly phase-separate upon osmotic cell volume change. Mol. Cell 2020, 79, 978–990.

(7) Marianelli, A. M.; Miller, B. M.; Keating, C. D. Impact of macromolecular crowding on RNA/spermine complex coacervation and oligonucleotide compartmentalization. Soft Matter 2018, 14, 368–378.

(8) Pullara, P.; Alshareedah, I.; Banerjee, P. R. Temperature-dependent reentrant phase transition of RNA-polycation mixtures. Soft Matter 2022, 18, 1342–1349.

(9) Hendrix, M.; Priestley, E. S.; Joyce, G. F.; Wong, C.-H. Direct Observation of Aminoglycoside–RNA Interactions by Surface Plasmon Resonance. J. Am. Chem. Soc. 1997, 119, 3641–3648.

(10) Kirk, S. R.; Tor, Y. tRNA^Phe^ Binds Aminoglycoside Antibiotics. Bioorg. Med. Chem. 1999, 7, 1979–1991.

(11) Wang, Y.; Rando, R. R. Specific Binding of Aminoglycoside Antibiotics to RNA. Chem. Biol. 1995, 2, 281–290.

(12) Vicens, Q.; Westhof, E. Crystal structure of geneticin bound to a bacterial 16S ribosomal RNA A site oligonucleotide. J. Mol. Biol. 2003, 326, 1175–1188.

(13) Demirci, H.; Murphy, F.; Murphy, E.; Gregory, S. T.; Dahlberg, A. E.; Jogl, G. A structural basis for streptomycin-induced misreading of the genetic code. Nat. Commun. 2013, 4, 1355.

(14) Jiang, L.; Majumdar, A.; Hu, W.; Jaishree, T. J.; Xu, W.; Patel, D. J. Saccharide-RNA recognition in a complex formed between neomycin B and an RNA aptamer. Structure 1999, 7, 817–827.

(15) Mikkelsen, N. E.; Johansson, K.; Virtanen, A.; Kirsebom, L. A. Aminoglycoside binding displaces a divalent metal ion in a tRNA–neomycin B complex. Nat. Struct. Biol. 2001, 8, 510–514.

(16) Wang, H.; Tor, Y. Electrostatic Interactions in RNA Aminoglycosides Binding. J. Am. Chem. Soc. 1997, 119, 8734–8735.

(17) Hermann, T.; Westhof, E. Docking of cationic antibiotics to negatively charged pockets in RNA folds. J. Med. Chem. 1999, 42, 1250–1261.

(18) Kopaczynska, M.; Schulz, A.; Fraczkowska, K.; Kraszewski, S.; Podbielska, H.; Fuhrhop, J. H. Selective condensation of DNA by aminoglycoside antibiotics. Eur. Biophys. J. 2015, 45, 287–299.

(19) Tripathi, V.; Bhadra, J.; Bhattacharya, S. Lipidated Aminoglycosides as New Tools that Repurpose Old Natural Antibiotics for New-Fangled Utility. Bioconjugate Chem. 2026, 37, 812–831.

(20) Kohanski, M. A.; Dwyer, D. J.; Hayete, B.; Lawrence, C. A.; Collins, J. J. A common mechanism of cellular death induced by bactericidal antibiotics. Cell 2007, 130, 797–810.

(21) Webster, C. M.; Shepherd, M. A mini-review: environmental and metabolic factors affecting aminoglycoside efficacy. World J. Microbiol. Biotechnol. 2022, 39, 7.

(22) Kalghatgi, S.; Spina, C. S.; Costello, J. C.; Liesa, M.; Morones-Ramirez, J. R.; Slo-movic, S.; Molina, A.; Shirihai, O. S.; Collins, J. J. Bactericidal antibiotics induce mitochondrial dysfunction and oxidative damage in mammalian cells. Sci. Transl. Med. 2013, 5, 192ra85.

(23) Karasawa, T.; Steyger, P. S. Intracellular mechanisms of aminoglycoside-induced cytotoxicity. Integr. Biol. 2011, 3, 879–886.

(24) Brangwynne, C. P.; Eckmann, C. R.; Courson, D. S.; Rybarska, A.; Hoege, C.; Gharakhani, J.; Jülicher, F.; Hyman, A. A. Germline P granules are liquid droplets that localize by controlled dissolution/condensation. Science 2009, 324, 1729–1732.

(25) Cheong, F. C.; Lee, S. Y.; Bais, S.; Khanal, S.; Saurabh, S. Holographic fingerprinting reveals oligomer-driven phase separation in Bovine Serum Albumin. Chem. Biomed. Imaging 2026,

(26) François, B.; Russell, R. J. M.; Murray, J. B.; Aboulela, F.; Masquida, B.; Vicens, Q.; Westhof, E. Crystal structures of complexes between aminoglycosides and decoding A site oligonucleotides: role of the number of rings and positive charges in the specific binding leading to miscoding. Nucleic Acids Res. 2005, 33, 5677–5690.

(27) Ennifar, E.; Paillart, J.-C.; Bodlenner, A.; Walter, P.; Weibel, J.-M.; Aubertin, A.-M.; Pale, P.; Dumas, P.; Marquet, R. Targeting the dimerization initiation site of HIV-1 RNA with aminoglycosides: from crystal to cell. Nucleic Acids Res. 2006, 34, 2328–2339.

(28) RCSB Protein Data Bank Molecular recognition of RNA by neomycin and a restricted neomycin derivative (PDB entry 2A04). Protein Data Bank, 2005; https://www.rcsb.org/structure/2A04.

(29) Bremer, H.; Dennis, P. P. Modulation of Chemical Composition and Other Parameters of the Cell at Different Exponential Growth Rates. EcoSal Plus 2008, 3.

(30) Bryan, L. E.; Kwan, S. Roles of ribosomal binding, membrane potential, and electron transport in bacterial uptake of streptomycin and gentamicin. Antimicrob. Agents Chemother. 1983, 23, 835–845.

(31) Sasazawa, M.; Tomares, D. T.; Childers, W. S.; Saurabh, S. Biomolecular condensates as stress sensors and modulators of bacterial signaling. PLoS Pathog. 2024, 20, e1012413.

(32) Michael, K.; Wang, H.; Tor, Y. Enhanced RNA Binding of Dimerized Aminoglycosides. Bioorg. Med. Chem. 1999, 7, 1361–1371.

(33) von Hofe, J.; Abacousnac, J.; Chen, M.; Sasazawa, M.; Javér Kristiansen, I.; Westrey, S.; Grier, D. G.; Saurabh, S. Multivalency controls the growth and dynamics of a biomolecular condensate. J. Am. Chem. Soc. 2025, 147, 25242–25253.

(34) Lee, S.-H.; Roichman, Y.; Yi, G.-R.; Kim, S.-H.; Yang, S.-M.; Van Blaaderen, A.; Van Oostrum, P.; Grier, D. G. Characterizing and tracking single colloidal particles with video holographic microscopy. Opt. Express 2007, 15, 18275–18282.

(35) Martin, C.; Altman, L. E.; Rawat, S.; Wang, A.; Grier, D. G.; Manoharan, V. N. In-line holographic microscopy with model-based analysis. Nat. Rev. Methods Primers 2022, 2, 83.

(36) Bohren, C. F.; Huffman, D. R. Absorption and Scattering of Light by Small Particles; Wiley Interscience: New York, 1983.

(37) Cornell, W. D.; Cieplak, P.; Bayly, C. I.; Gould, I. R.; Merz, K. M.; Ferguson, D. M.; Spellmeyer, D. C.; Fox, T.; Caldwell, J. W.; Kollman, P. A. A Second Generation Force Field for the Simulation of Proteins, Nucleic Acids, and Organic Molecules. J. Am. Chem. Soc. 1995, 117, 5179–5197.

(38) Pérez, A.; Marchán, I.; Svozil, D.; Sponer, J.; Cheatham, T. E., III; Laughton, C. A.; Orozco, M. Refinement of the AMBER Force Field for Nucleic Acids: Improving the Description of *α*/*γ* Conformers. Biophys. J. 2007, 92, 3817–3829.

(39) Zgarbová, M.; Otyepka, M.; Sponer, J.; Mládek, A.; Banaš, P.; Cheatham, T. E., III; Jurečka, P. Refinement of the Cornell et al. Nucleic Acids Force Field Based on Reference Quantum Chemical Calculations of Glycosidic Torsion Profiles. J. Chem. Theory Comput. 2011, 7, 2886–2902.

(40) Wang, J.; Wolf, R. M.; Caldwell, J. W.; Kollman, P. A.; Case, D. A. Development and Testing of a General Amber Force Field. J. Comput. Chem. 2004, 25, 1157–1174.

(41) Jakalian, A.; Jack, D. B.; Bayly, C. I. Fast, Efficient Generation of High-Quality Atomic Charges. AM1-BCC Model: II. Parameterization and Validation. J. Comput. Chem. 2002, 23, 1623–1641.

(42) Jorgensen, W. L.; Chandrasekhar, J.; Madura, J. D.; Impey, R. W.; Klein, M. L. Comparison of Simple Potential Functions for Simulating Liquid Water. J. Chem. Phys. 1983, 79, 926–935.

(43) Joung, I. S.; Cheatham, T. E., III Determination of Alkali and Halide Monovalent Ion Parameters for Use in Explicitly Solvated Biomolecular Simulations. J. Phys. Chem. B 2008, 112, 9020–9041.

(44) Darden, T.; York, D.; Pedersen, L. Particle Mesh Ewald: An *N* · log(*N*) Method for Ewald Sums in Large Systems. J. Chem. Phys. 1993, 98, 10089–10092.

(45) Roe, D. R.; Cheatham, T. E., III PTRAJ and CPPTRAJ: Software for Processing and Analysis of Molecular Dynamics Trajectory Data. J. Chem. Theory Comput. 2013, 9, 3084–3095.

(46) Alkhzem, A. H.; Woodman, T. J.; Blagbrough, I. S. Multinuclear Nuclear Magnetic Resonance Spectroscopy Is Used to Determine Rapidly and Accurately the Individual p*K*_a_ Values of 2-Deoxystreptamine, Neamine, Neomycin, Paromomycin, and Streptomycin. ACS Omega 2021, 6, 2824–2835.

